# Construction of a constitutively active type III secretion system for heterologous protein secretion

**DOI:** 10.1101/2022.02.24.481905

**Authors:** Julie Ming Liang, Lisa Ann Burdette, Han Teng Wong, Danielle Tullman-Ercek

## Abstract

Proteins comprise a multibillion-dollar industry in enzymes and therapeutics, but bacterial protein production can be costly and inefficient. Proteins of interest (POIs) must be extracted from lysed cells, purified, and resolubilized. The *Salmonella* pathogenicity island 1 type III secretion system (T3SS) is a protein secretion complex in *Salmonella* that has been engineered to secrete heterologous proteins and addresses the problems associated with bacterial protein production. However, the current best practices method of T3SS pathway activation for secretion is not ideal for industrial scaleup. Previously, the T3SS was activated by plasmid-based overexpression of the T3SS transcriptional regulator, *hilA*, which requires the addition of a small molecule inducer to the culture media and adds significant cost to the production media. Plasmid-based expression is also subject to instability in large-scale fermentation. Here, we show that we can constitutively activate the T3SS by modulating the upstream transcriptional regulator, *hilD*, either through knocking out *hilE*, a repressor of HilD, or by adding transcriptional fusions to *hilD*. Finally, we combine the two most promising genomic modifications to build a constitutively active T3SS capable of secreting a range of heterologous proteins at titers comparable to those reported with synthetic induction of *hilA*. These improvements further our goal of making an industrially competitive protein production strain that reduces the challenges associated with plasmid induction and maintenance.

**Importance:** Proteins are used in our everyday lives as therapeutics (insulin), industrial enzymes (laundry detergent), and bio-based materials (spider silk). Current industrial protein production in bacteria is costly because it requires purification of the target protein from the other proteins inside the cell. We solve this problem by engineering the *Salmonella* pathogenicity island 1 type III secretion system (T3SS) to export the target protein into the cell growth media. This makes the protein purification process more efficient and cheaper. However, this system currently requires an expensive inducer reagent to activate it which significantly increases the cost of the production media. We show here the creation of a constitutively active T3SS, meaning the T3SS pathway is always on. In doing so, we successfully created a *Salmonella* strain that eliminates the need for the inducer reagent and exports proteins at levels comparable to the inducer-activated system, ultimately reducing the cost of T3SS protein production.

## Introduction

Recombinant proteins are ubiquitous in modern society and are used in diverse applications as pharmaceutical agents, research tools, active ingredients in consumer products, and catalysts in industrial processes. Bacteria are good hosts for producing these recombinant proteins given their genetic tractability, fast growth, and high yield. Traditionally, recombinant protein expression in gram-negative bacteria is intracellular, and the cells must be lysed to recover the protein of interest (POI). This process can be time-consuming and resource intensive as recovering a pure product from the cellular contents can require several downstream purification steps. Further, recombinant proteins often accumulate in insoluble aggregates known as inclusion bodies, which can increase initial purity but also requires significant process optimization to resolubilize and refold the product.^1–3^ Finally, intracellular protein expression is not ideal for proteins that are cytotoxic, prone to proteolysis, or self-assembling. Secreting the protein product outside of the bacterial cell retains the benefits of traditional bacterial protein expression while addressing many of its challenges.^4–6^

Gram-negative bacteria possess five secretion systems that can be engineered to secrete recombinant proteins.^7^ The type III secretion system (T3SS) is a transmembrane protein complex that is attractive for recombinant protein production because of its ability to translocate proteins from the cytoplasm to the culture media in a single step.^8, 9^ The T3SS is nonessential for viability, which enables repurposing it solely for heterologous protein secretion. In particular, the *Salmonella* pathogenicity island 1 T3SS (SPI-1 T3SS) of *Salmonella enterica* serovar Typhimurium has been engineered to secrete a variety of industrially relevant proteins including enzymes, biomaterials, and antibody fragments.^5, 6, 9, 10^

Secretion via the SPI-1 T3SS requires activation of the system; it is not constitutively expressed. The native SPI-1 T3SS is activated by environmental cues, which initiate a feed-forward loop consisting of the transcription factors HilD, HilC, and RtsA.^11^ This feed-forward loop in turn activates the SPI-1 T3SS regulator HilA, which then activates transcription of the operons containing genes that code for T3SS structural and regulatory proteins.^12, 13^ HilD is the upstream regulator to *hilA*, acts with HilC and RtsA to regulate expression of *hilA*, and controls the commitment step of *Salmonella* to the SPI-1 T3SS pathway.^11, 14, 15^ The SPI-1 T3SS is commonly activated in a laboratory environment by growing *S. enterica* in high salt, low aeration conditions.^9, 16^ However, these conditions limit achievable cell densities and they result in an active T3SS in only ∼33% of the population.^17^

Our lab previously designed a synthetic activation strategy to overcome these limitations.^10^ Plasmid-based overexpression of *hilA* allowed for T3SS activation in standard laboratory growth conditions and media, increasing both the maximum cell density and the percentage of the population expressing the T3SS (>90%). These improvements increased protein secretion titers >10-fold. However, plasmid-based overexpression of *hilA* is not ideal for industrial purposes because it requires the addition of an expensive inducer molecule and adds another plasmid to our system, as the secreted protein is expressed from a plasmid. The inducer and antibiotic required to maintain the second plasmid make up 21% (industrial pricing) - 28% (academic pricing) of the total media cost (Table 1). Plasmids also impose a significant metabolic burden, and they are prone to instability in industrial-scale cultures.^18, 19^ Therefore, realization of the SPI-1 T3SS as a recombinant protein production platform would benefit immensely from a regulation strategy that results in similar activation levels and titers without requiring the *hilA* activation plasmid or the addition of an inducer. However, this goal is complicated by the fact that the SPI-1 T3SS is sensitive to environmental inputs, a property we recently leveraged to design an optimized growth medium for protein secretion.^20^ Manipulating expression of one of the members of the feed-forward loop that integrates environmental signals upstream of *hilA* could help achieve these goals while circumventing sensitivities to environmental inputs.

**Table 1.** Reagent costs used to calculate percentage cost of inducer. “Cost (US$/kg)” is the cost of one kilogram of that reagent and “Cost (US$/L)” is the cost of using that reagent in 1 liter of production media.

| Reagent | Media g/L | Academic cost |  | Industrial cost |  |
| --- | --- | --- | --- | --- | --- |
| | | Cost (US\$/kg) | Cost (US\$/L) | Cost (US\$/kg) | Cost (US\$/L) |
| Tryptone | 10 | 75.58 | 0.7558 | 27.4<sup>a</sup> | 0.274 |
| Yeast Extract | 5 | 265.8 | 1.329 | 1.7<sup>b</sup> | 0.0085 |
| NaCl | 5 | 12.78 | 0.0639 | 0.08<sup>c</sup> | 0.0004 |
| Kanamycin | 0.5 | 3934 | 1.967 | 1<sup>b</sup> | 0.0005 |
| Chloramphenicol | 0.34 | 1037.2 | 0.3526 | 43<sup>d</sup> | 0.01462 |
| IPTG | 0.1 | 12700 | 1.27 | 601<sup>b</sup> | 0.0601 |
| Estimated media cost (with inducer) |  |  | 5.7383 |  | 0.35812 |
| Estimated media cost (without inducer and antibiotic) |  |  | 4.1157 |  | 0.2834 |

<sup>a</sup>Li, ZN *et al.* doi.org/10.1007/s13399-020-00800-3.
<sup>b</sup>Cardoso, VM *et al.* doi.org/10.1016/j.btre.2020.e00441.
<sup>c</sup>Wu, N *et al.* doi.org/10.1016/j.jclepro.2020.124939.
<sup>d</sup>Molbase.

Of the three members of the feed-forward loop that controls SPI-1 activation, HilD has a dominant role.^11, 14, 21, 22^ We explored strategies to change HilD levels via genomic modifications as HilD expression and function are controlled at the transcriptional and post-translational levels.^21, 23, 24^ Controlling transcriptional regulator levels by genome editing eliminates the need for an activation plasmid and inducer for T3SS activation. In this study, we confirm that overexpression of *hilD* can activate the SPI-1 T3SS pathway for heterologous protein secretion and match secretion titers achieved with *hilA* overexpression. We constructed a strain with a constitutively active SPI-1 T3SS by deleting *hilE*, which negatively regulates HilD by blocking its ability to bind DNA,^24^ and by transcriptionally fusing reporter proteins downstream of *hilD*. Transcriptional activity assays showed that these genomic modifications resulted in constitutive activation of the T3SS in >90% of the population. Finally, we combined the *hilE* deletion and a *hilD* transcriptional fusion to create a strain that had a constitutively active T3SS in >99% of the population. The engineered strain expresses and secretes recombinant proteins at levels equivalent to the current plasmid-based *hilA* overexpression strategy. This new strain lowers the overall cost of fermentation by reducing the number of required media components (inducer and antibiotic for the second plasmid), ultimately furthering the development of a T3SS-based protein production platform.

## Materials and Methods

### Strains and Genomic Modifications

Strains and primers used to construct the strains in this study are listed in Table 2 **and** 3. The parent strain used to construct all the strains described in this work, ASTE13, is a high-secreting derivative of the LT2 strain of *S. enterica* serovar Typhimurium.^20^ Genomic modifications for this study were created using the λ Red recombineering method developed by the Court lab.^25^ This method uses a counter-selectable marker, *catG-sacB*, to scarlessly incorporate changes into a bacterial genome. Briefly, the *catG-sacB* cassette is incorporated at the correct site and selected for by chloramphenicol resistance. Then, the cassette is replaced with the target DNA construct and selected by growth on sucrose plates. The *catG-sacB* cassette was amplified from the *Escherichia coli* TUC01 genome with primers containing 40 bp of homology 5’ or 3’ of the appropriate genetic locus strains. Coding sequences were amplified as double-stranded DNA fragments with the same 40 bp homology 5’ and 3’ as the *catG-sacB* cassette to facilitate recombination at the correct site. Knockouts were performed by introducing a 70 bp oligo composed of the 35 bp of homology upstream and downstream of the deletion site stitched together. Double-stranded DNA used to introduce cassettes for recombineering was amplified using Phusion polymerase (New England Biolabs). Successful recombinants were confirmed by Sanger sequencing (Quintara Biosciences) of a colony PCR product of the appropriate locus. Colony PCR was performed using GoTaq polymerase (Promega). The Δ*hilE hilD::mCherry* was constructed by incorporating *hilD::mCherry* construct into the Δ*hilE* strain.

**Table 2.** Strains used in this study.

| Strain Name | Description | Reference |
| --- | --- | --- |
| ASTE13 | LT2-derived lab strain similar to DW01 | Burdette et al<sup>a</sup> |
| ASTE13 $\Delta invA$ | *invA* knockout. InvA is required for functional SPI-1 T3SS secretion | This study |
| ASTE13 $\Delta hilE$ | *hilE* knockout. HilE is a SPI-1 T3SS repressor | This study |
| ASTE13 *hilD::GFP* | *GFPmut2* is inserted downstream of the *hilD* stop codon | This study |
| ASTE13 *hilD::mCherry* | *mCherry* is inserted downstream of the *hilD* stop codon | This study |
| ASTE13 *hilD::halotag* | *halotag* is inserted downstream of the *hilD* stop codon | This study |
| ASTE13 *hilD::MBP* | Maltose binding protein is inserted downstream of the *hilD* stop codon | This study |
| ASTE13 $\Delta hilE$ *hilD::mCherry* | *hilE* knockout and *mCherry* is inserted downstream of the *hilD* stop codon | This study |

<sup>a</sup>Burdette, LA *et al.* doi.org/10.1186/s12934-021-01536-z.

### Plasmids and Plasmid Construction

Plasmids and primers used to construct the plasmids in this study are listed in Table 3 **and** 4. The *P_lacUV5_ hilD* plasmid was constructed by amplifying the *hilD* coding sequence including 20 bp upstream of the start codon from ASTE13 and cloned by Gibson assembly into the *P_lacUV5_ hilA* plasmid without the *hilA* gene.^10^ The *P_prg_* GFP plasmid was previously unpublished and constructed by Dr. Kevin J. Metcalf using the same method as the *P_sic_* GFP and *P_inv_* GFP plasmids in Metcalf et al. The GFP variant is GFPmut2.^10^

**Table 3.**
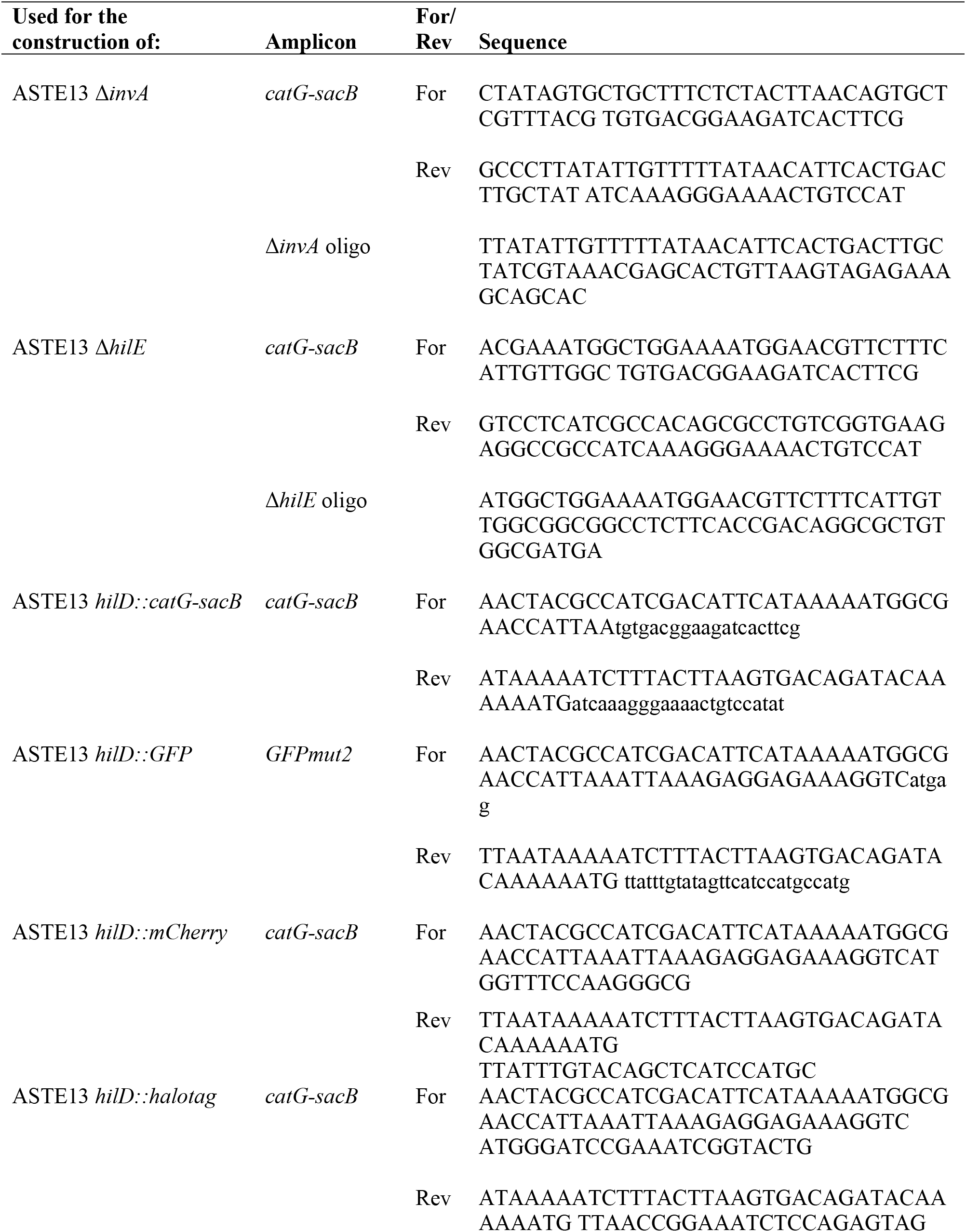
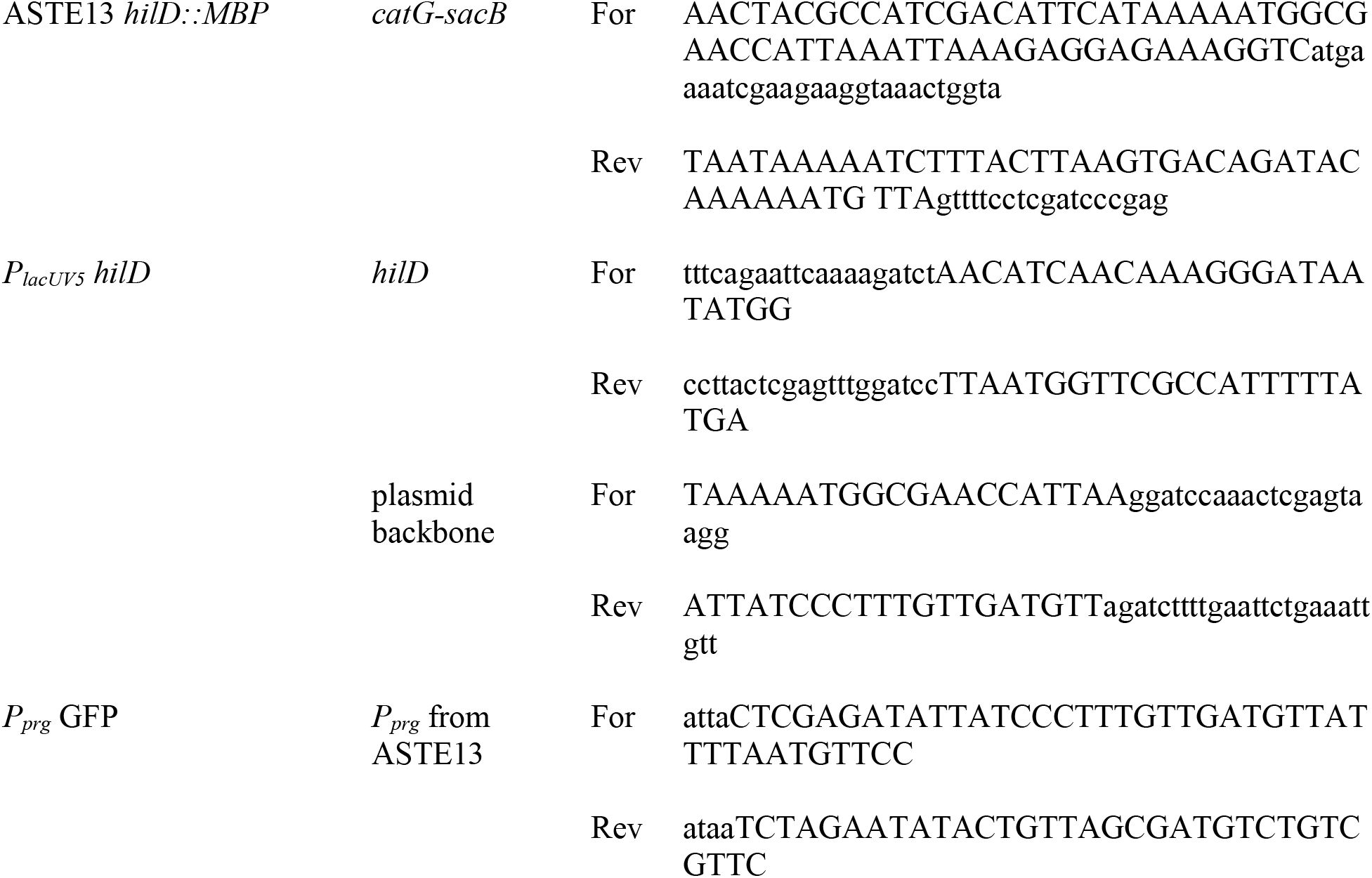
Primers used in this study.

**Table 4.** Plasmids used in this study.

| Plasmid Name | ORF Description | Antibiotic Resistance | Origin of Replication | Reference |
| --- | --- | --- | --- | --- |
| *P<sub>lacUV5</sub> hilA* | *hilA* | kan | colE1 | Metcalf et al.<sup>a</sup> |
| *P<sub>lacUV5</sub> hilD* | *hilD* | kan | colE1 | This study |
| *P<sub>sic</sub> SptP-DH* | *sicP sptP-DH-2xF-6xH* | cam | p15a | Widmaier et al.<sup>b</sup> |
| *P<sub>tet</sub> SptP-DH* | *sicP sptP-DH-2xF-6xH* | cam | p15a | Metcalf et al.<sup>c</sup> |
| *P<sub>sic</sub> GFP* | *GFPmut2* | cam | p15a | Metcalf et al.<sup>c</sup> |
| *P<sub>prg</sub> GFP* | *GFPmut2* | cam | p15a | This study |
| *P<sub>inv</sub> GFP* | *GFPmut2* | cam | p15a | Metcalf et al.<sup>c</sup> |
| *P<sub>sic</sub> SopB-DH* | *sigE sopB-DH-2xF-6xH* | cam | p15a | Widmaier et al.<sup>b</sup> |
| *P<sub>sic</sub> SptP-phoA* | *sicP sptP-phoA-2xF-6xH* | cam | p15a | Metcalf et al.<sup>a</sup> |
| *P<sub>sic</sub> SptP-hGH* | *sicP sptP-hGH-2xF-6xH* | cam | p15a | Burdette et al.<sup>c</sup> |

<sup>a</sup>Metcalf, KJ *et al.* doi.org/10.1128/AEM.01330-14. <sup>b</sup>Widmaier, DM *et al.* doi.org/10.1038/msb.2009.62. <sup>c</sup>Burdette, LA *et al.* doi.org/10.1186/s12934-021-01536-z.

### Secretion Growth Conditions

Cells were transformed with the required plasmids via electroporation. Secretion experiments (e.g., using cells harboring *P_sic_* SptP-DH, *P_sic_* SopB-DH, *P_sic_* SptP-rhGH, and *P_sic_* SptP-PhoA) were started from a single colony grown overnight in the lysogeny broth (LB) Lennox formulation with appropriate antibiotics (34 µg/mL chloramphenicol and/or 50 µg/mL kanamycin) at 37°C and 225 rpm in 24-well blocks (Axygen) sealed with AeraSeal film to allow for air exchange. Cultures for secretion and regulation experiments were subcultured from overnights to 0.05 OD_600_ in 5 mL LB-Lennox supplemented with appropriate antibiotics in 24-well blocks. When appropriate, 100 µg/mL isopropyl β-d-1-thiogalactopyranoside (IPTG) was added at the time of subculture to induce *P_lacUV5_ hilA* or *P_lacUV5_ hilD* expression. Cultures were grown for 8 hours at 37°C and 225 rpm to facilitate protein secretion. At the time of harvest, whole culture lysate (WCL) samples for sodium dodecyl sulphate–polyacrylamide gel electrophoresis (SDS-PAGE) were collected by adding 20 µL of final cell culture to 40 µL of 4X Laemmli buffer. Cells were pelleted in the 24-well blocks at 4000 x *g* for 10 minutes to isolate the secreted fraction. SDS-PAGE samples for the secreted fraction were prepared by adding 40 µL of supernatant to 16 µL of 4X Laemmli buffer. All SDS-PAGE samples were boiled at 95°C for 5 minutes after mixing. Statistical significance was determined using Welch’s unequal variances t-test.

### Optimal Induction Timing Secretion Growth Conditions

Strains were transformed with *P_tet_* SptP-DH and cultured overnight as described above in the previous section, “Secretion Growth Conditions”. The next morning, strains were subcultured from the overnights to 0.05 OD_600_ in 5 mL LB-Lennox supplemented with the appropriate antibiotics (34 µg/mL chloramphenicol and 50 µg/mL kanamycin) in 24-well blocks. Strains were grown at 225 rpm and 37°C. Each overnight strain culture was subcultured into 6 wells, and each well was induced with anhydrotetracycline (aTc, 100 ng/mL) every hour from 0-5 hours after subculture to activate the *P_tet_* DH plasmid. Whole culture lysate and secreted fraction samples were collected after 8 hours, prepared, and analyzed as described in the previous section.

### Transcriptional Activity Assay and Flow Cytometry

Each indicated strain was transformed via electroporation with a plasmid encoding a designated SPI-1 promoter (*P_sic_*, *P_prg_*, or *P_inv_*) driving the expression of GFPmut2. The strains were then grown, diluted, and induced as described above in the previous section, “Secretion Growth Conditions”, except the overnight cultures were supplemented with 0.4% w/v glucose to repress expression of the fluorescent reporter. Samples were taken every hour for 8 hours and diluted to an OD_600_ of approximately 0.03 in phosphate-buffered saline (PBS) with 2 mg/mL of kanamycin sulfate in round bottom plates (Greiner Bio-One #650101). Samples were run on an Attune NxT Flow Cytometer via the autosampler on the same day and at least 20,000 events per sample were captured. Forward scatter, side scatter, and fluorescence on the BL1 channel (ex: 488nm, em: 530/30nm) were measured (voltages below). Samples were gated for cells and singlets. The T3SS-active population was determined by gating the events above the fluorescence of the wildtype (WT, no activation plasmid) population with the GFP transcriptional reporter. Data was processed with FlowJo v10.5.3 (TreeStar, Inc.) and visualized with R and the ggplot2 package.^26^ Arbitrary units were converted to Molecules of Equivalent Fluorescence (MEF) using a calibration curve.

The calibration curve was constructed using BD Sphero Rainbow Calibration Particles (B559123) according to manufacturer’s guidelines.

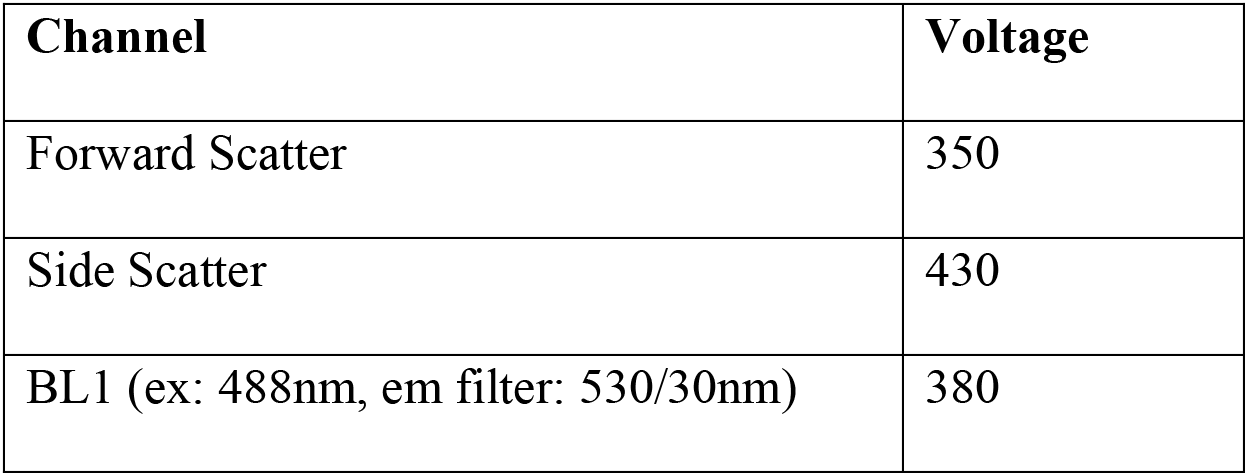

### SDS-PAGE and Western Blotting

Whole culture lysate (WCL) and secreted fraction (SF) samples were separated by size on 12.5% SDS-PAGE gels run at 150V for 60 minutes using the Mini-PROTEAN Tetra Cell system (Bio-Rad). The proteins were transferred to a polyvinylidene fluoride membrane (PVDF, Millipore) using the Trans-Blot SD Semi-Dry Transfer Cell (Bio-Rad). Membranes were probed with mouse anti-FLAG M2 antibody (Millipore Sigma F3165, 1:6,666 dilution) according to manufacturer’s instructions. The anti-FLAG antibody was subsequently detected with goat anti-mouse IgG (H+L) conjugated to horseradish peroxidase (HRP) to facilitate chemiluminescent detection (Invitrogen 32430, 1:1,000 dilution). Chemiluminescent signal was produced with Pierce ECL substrate (Thermo Scientific), and blots were imaged using a ChemiDoc XRS+ (Bio-Rad).

Relative secretion titer was quantified by normalizing densitometry values calculated using Image Lab software (v5.2.1, Bio-Rad). Densitometry values were normalized by first dividing by the OD_600_ measurement, and then dividing by the appropriate OD-normalized control strain, typically ASTE13 WT carrying *P_lacUV5_* HilA and *P_sic_* SptP-DH-2xFLAG-6xHis present on each blot.

To detect GroEL, anti-FLAG blots were stripped using a mild stripping protocol modified from Abcam. The membranes were incubated in mild stripping buffer (15 g/L glycine, 1 g/L SDS, 1% v/v Tween 20, pH 2.2) for two periods of 8 minutes, washed twice in ddH_2_O for 10 minutes, and then washed twice in TBST for 5 minutes. The stripped blot was blocked in 5% w/v nonfat milk in TBST. The membrane was probed with rabbit anti-GroEL according to manufacturer’s instructions (Sigma Aldrich, 1:10,000 dilution). Membranes were subsequently probed with goat anti-rabbit IgG (H+L) conjugated to HRP according to manufacturer’s instructions (Thermo Fisher, 1:1,000 dilution). Bands were detected with SuperSignal West Pico Plus substrate (Thermo Fisher) and a ChemiDoc XRS+ imaging system.

### Growth Media Cost Analysis

The growth media cost analysis was calculated using academic and industry pricing^27–30^ for reagents used in this study and described in Table 1. The percentage cost of IPTG was calculated by comparing the total cost of the production media with and without inducer and the necessary antibiotic to sustain that plasmid. The production media is LB-L supplemented with 34 µg/mL chloramphenicol and 50 µg/mL kanamycin and the components of LB-L are 10g/L tryptone, 5 g/L yeast extract, and 5 g/L sodium chloride. “Cost (US$/kg)” is the cost of one kilogram of that reagent and “Cost (US$/L)” is the cost of using that reagent in 1 liter of media.

## Results

### Inducible overexpression of T3SS master regulator HilD produces comparable secretion titers to *hilA* overexpression

Overexpression of *hilD* is a common strategy for synthetic activation of the SPI-1 T3SS for regulatory studies,^31, 32^ but *hilD* overexpression has not yet been used to activate the T3SS for heterologous protein secretion. Though our overarching goal is to replace plasmid-based T3SS activation with genomic modifications that cause constitutive T3SS expression, we first sought to compare plasmid-based *hilD* overexpression to the already optimized *hilA* overexpression strategy to validate the *hilD* overexpression approach in this context. To this end, we constructed an isopropyl β-d-1-thiogalactopyranoside (IPTG) inducible low-copy plasmid, *P_lacUV5_ hilD*, and co-transformed it with the export plasmid. The export plasmid has been used previously and encodes for a model secreted protein, the DH domain of the human protein intersectin-1L, fused to the T3SS secretion tag SptP.^9, 33^ SptP-DH-2xFLAG-6xHis (hereafter, SptP-DH) was expressed under the control of the T3SS promoter *P_sic_*. The *sic* promoter drives expression of native secreted T3SS proteins and is activated by a cascade following T3SS activation. Otherwise, expression of the POI would require an additional inducer. By using the *sic* promoter to drive expression of the POI, we can express both the T3SS machinery and the protein of interest (POI) following one activation event rather than two. To find the optimum induction level for *hilD* overexpression, the *P_lacUV5_ hilD* plasmid was induced by IPTG at concentrations ranging from 0 µM to 1000 µM. We used semi-quantitative western blotting to compare the resulting SptP-DH secretion titers to the current standard T3SS activation method, *hilA* overexpression.^10^

Induction of *P_lacUV5_ hilD* resulted in SptP-DH secretion titers that increased with IPTG concentration, confirming that overexpression of *hilD* is sufficient for activating the SPI-1 T3SS for heterologous protein secretion (Figure 1). Secretion titer increases with IPTG concentration until 500 µM. IPTG concentrations above 50 µM produce secretion titers equal to or higher than optimized *hilA* overexpression. A similar trend is apparent in the SptP-DH expression data (**Supplemental Figure S1**). This suggests that secretion titer increases with expression level when the T3SS is induced by *hilD* overexpression (R^2^ = 0.72, **Supplemental Figure S2**). The secretion titer resulting from *hilD* overexpression at 100 µM IPTG is not significantly different from *hilA* overexpression at the same level (p-value = 0.482). Notably, a growth defect is observed when *hilD* overexpression is induced with >100 µM IPTG (**Supplemental Table S1**). For these reasons, we used an IPTG concentration of 100 µM for subsequent analyses with *P_lacUV5_ hilD* to achieve a balance between growth and secretion titer.

**Figure 1:**
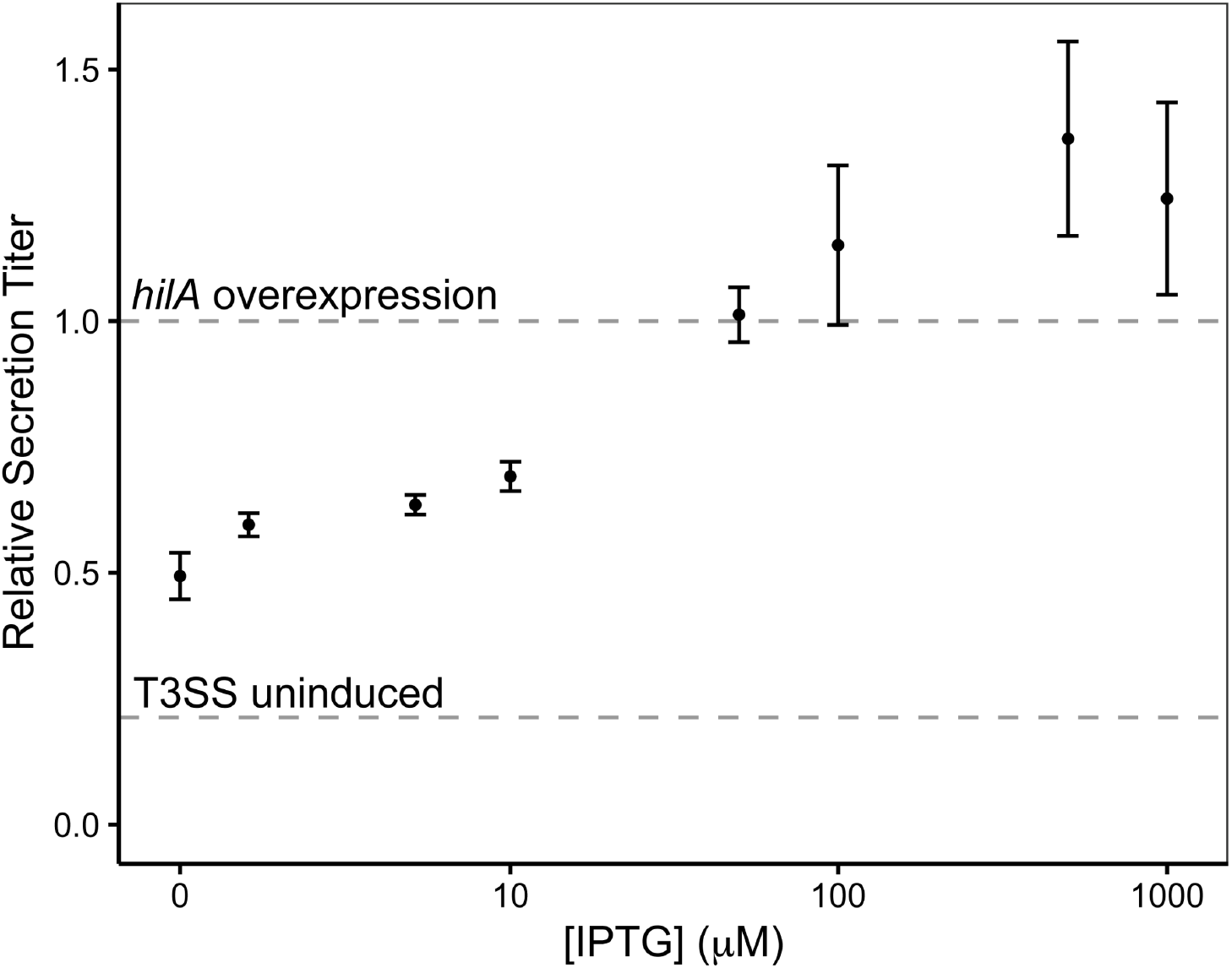
Effect of *hilD* overexpression at varying induction levels on SptP-DH secretion titer. A strain harboring an export plasmid encoding SptP-DH and an activation plasmid with *hilD* under the control of a *lac* promoter was induced with IPTG at concentrations ranging from 0 µM to 1000 µM. Relative secretion titers were measured by western blot of secreted fraction samples and probed against a FLAG tag on the SptP-DH construct. Densitometry results were normalized by OD_600_ and *hilA* overexpression at 100 μM IPTG per replicate (top dashed reference line). The bottom dashed reference line indicates relative SptP-DH secretion titer in a strain without any synthetic induction plasmid. Data represents three replicates and error bars represent standard error.

To ensure that secretion was specific to the SPI-1 T3SS, we next tested *hilD* overexpression in an *invA* deletion strain (Δ*invA*). The Δ*invA* strain is a T3SS secretion-incompetent strain that does not produce functional SPI-1 T3SS apparatuses.^9, 34^ Overexpression of *hilD* did not result in detectable SptP-DH secretion in the Δ*invA* strain (**Supplemental Figure S3**), indicating that secretion by *hilD* overexpression occurs through the SPI-1 T3SS rather than a different T3SS (flagellar or SPI-2). As an additional control, we verified that the presence of SptP-DH in the secreted fraction was not due to cell lysis by western blotting against GroEL, an intracellular chaperone protein.^9, 10, 35^ GroEL levels were similar among strains overexpressing *hilD*, overexpressing *hilA*, or lacking a synthetic overexpression plasmid (WT), indicating that *hilD* overexpression did not cause an increase in cell lysis (**Supplemental Figure S4**). Together these results indicate that *hilD* overexpression induced T3SS-dependent protein secretion to similar levels to that of *hilA* overexpression.

### *Sic* promoter-driven POI expression results in equivalent or higher secretion titers

Since *hilD* overexpression is a new, uncharacterized method of activation for T3SS-based heterologous protein secretion, we sought to determine whether the *sic* promoter is still the optimal method for activating expression of the model POI, SptP-DH. As a reminder, the previous strategy to activate the T3SS with *hilA* overexpression used the *sic* promoter to drive expression of the POI. This was advantageous because the *sic* promoter is activated following the activation of the SPI-1 T3SS and enabled automatic expression of the POI without the need for two inducers (one for the *hilA* activation plasmid and one for the POI). To evaluate the continued use of the *sic* promoter, we examined expression and secretion titers using the *tet* promoter, activated by the small molecule anhydrotetracycline (aTc), and compared them to those obtained using the native *sic* promoter. First, we determined the optimal induction time by adding aTc (100 ng/mL) to the culture media every hour from 0-5 hours after inducing the *hilA* or *hilD* overexpression plasmids. We measured the resulting secretion titers and expression using semi-quantitative western blotting (**Supplemental Figure S5 & S6**) and identified the induction time that resulted in the highest secretion titer for *hilA* overexpression (3 hours) and *hilD* overexpression (1 hour). We then compared the secretion titer and expression of SptP-DH between the inducible promoter, *P_tet_*, induced at the time that gave highest titer, and the native promoter, *P_sic_*.

For *hilD* overexpression, the secretion titer of SptP-DH expressed using *P_tet_* was not significantly different from the secretion titer of SptP-DH expressed using the *sic* promoter (Figure 2). This data shows that we can effectively drive POI expression using the *P_sic_* promoter when the T3SS is activated by *hilD* overexpression, which is compatible with our goal of developing an inducer-free platform for protein secretion with the T3SS.

**Figure 2:**
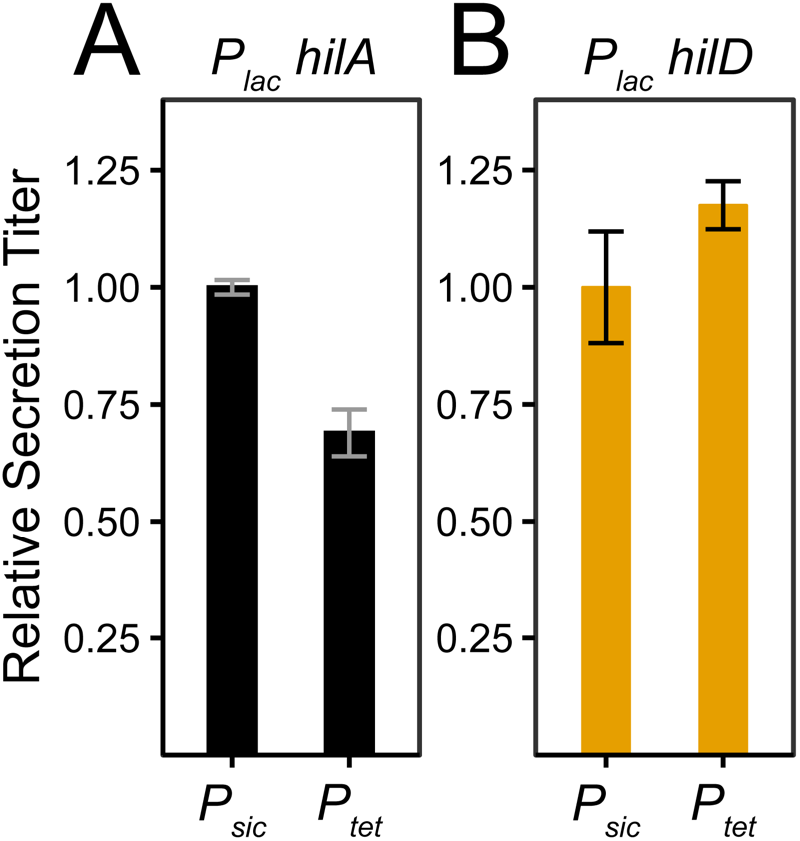
Comparison of secretion titers when SptP-DH is expressed from the native *sic* promoter versus the *tet* promoter in two contexts. **(A)** Comparison of strains containing the activation plasmid *P_lac_ hilA* and an export plasmid encoding either *P_sic_* SptP-DH or *P_tet_* SptP-DH. **(B)** Comparison of strains containing the activation plasmid *P_lac_ hilD* and an export plasmid encoding either *P_sic_* SptP-DH or *P_tet_* SptP-DH. *P_tet_* DH is induced with aTc (100 ng/mL) at the optimal induction time for protein secretion (at 3 hours for *hilA* and 1 hour for *hilD* overexpression). *P_lac_ hilA* and *P_lac_ hilD* were induced at 100 μM IPTG. Relative secretion titer was measured by western blot of secreted fraction samples and probed against a FLAG tag on the SptP-DH construct. The titer was normalized by the average densitometry of the *P_sic_* DH densitometry for each activation method. Data represents three replicates and error bars represent standard error.

### Overexpression of *hilD* causes early activation and deactivation of T3SS activity

Although the secretion titers between *hilA* and *hilD* activation are similar, the two methods may have different impacts on T3SS transcriptional activity. This is further supported by the fact that POI expression levels between *hilA* and *hilD* overexpression were different even though both overexpression strategies resulted in similar secretion titers (Figure 2 and **Supplemental Figure S7**). Thus, we sought to characterize differences in T3SS transcriptional activity between the two activation strategies to better understand the differences so that we could further engineer this system. To this end, we evaluated the impact of *hilD* overexpression on the transcriptional activity of several key SPI-1 operon promoters (*prg*, *inv*, and *sic*) that drive expression of the physical T3SS apparatuses and secreted proteins. The *prg* and *inv* operons encode for the structural components of the T3SS apparatus, while the *sic* operon encodes for the secreted effectors. The *sic* operon is under control of the *P_sic_* promoter, which also controls secreted POI expression in the export plasmid used in this study. We measured transcriptional activity using a flow cytometry-based assay in which GFP expression is controlled by these SPI-1 T3SS promoters. Thus, GFP fluorescence provides a quantifiable readout of transcriptional activity. We determined the effect of *hilD* overexpression on both the percentage of cells with activated T3SS promoters (“T3SS active”) in the population and the average activity level as indicated by the geometric mean fluorescence intensity (MFI) of the active population.

Previously, *hilA* overexpression increased secretion titer by increasing both the number of cells in the population with T3SS activity and the level of T3SS activity within a given cell.^10^ Synthetic overexpression of *hilD* produces a similar result for *P_sic_* activity, in which >90% of the population is T3SS active. In contrast, the strain lacking an induction plasmid (WT) peaked at 65% of the population showing *sic* promoter activity (Figure 3). MFI, which represents the level of T3SS activation, reaches similar levels for *hilA* and *hilD* overexpression. Interestingly, although *hilA* and *hilD* overexpression results in a similar percent activated population and MFI at the end of 8 hours, the two overexpression strategies had different activation times for the *sic* operon. We observe increased activity after 1 hour post-induction via overexpression of *hilD*, compared to after 3 hours via *hilA* overexpression and between 3 and 4 hours for the WT strain lacking a synthetic overexpression plasmid. This correlates with the optimal induction time for expressing the POI that we determined earlier (**Supplemental Figure S5**). The temporal difference between *hilA* overexpression and the WT strain lacking a synthetic overexpression plasmid was documented previously.^10^ However, the temporal variability between *hilA* and *hilD* overexpression is surprising because *hilD* is not known to interact directly with *P_sic_* and is upstream of *hilA* in the T3SS regulatory cascade. We therefore had expected the *hilD* overexpression strategy to take longer to activate *P_sic_* because it must first activate *hilA*.

**Figure 3:**
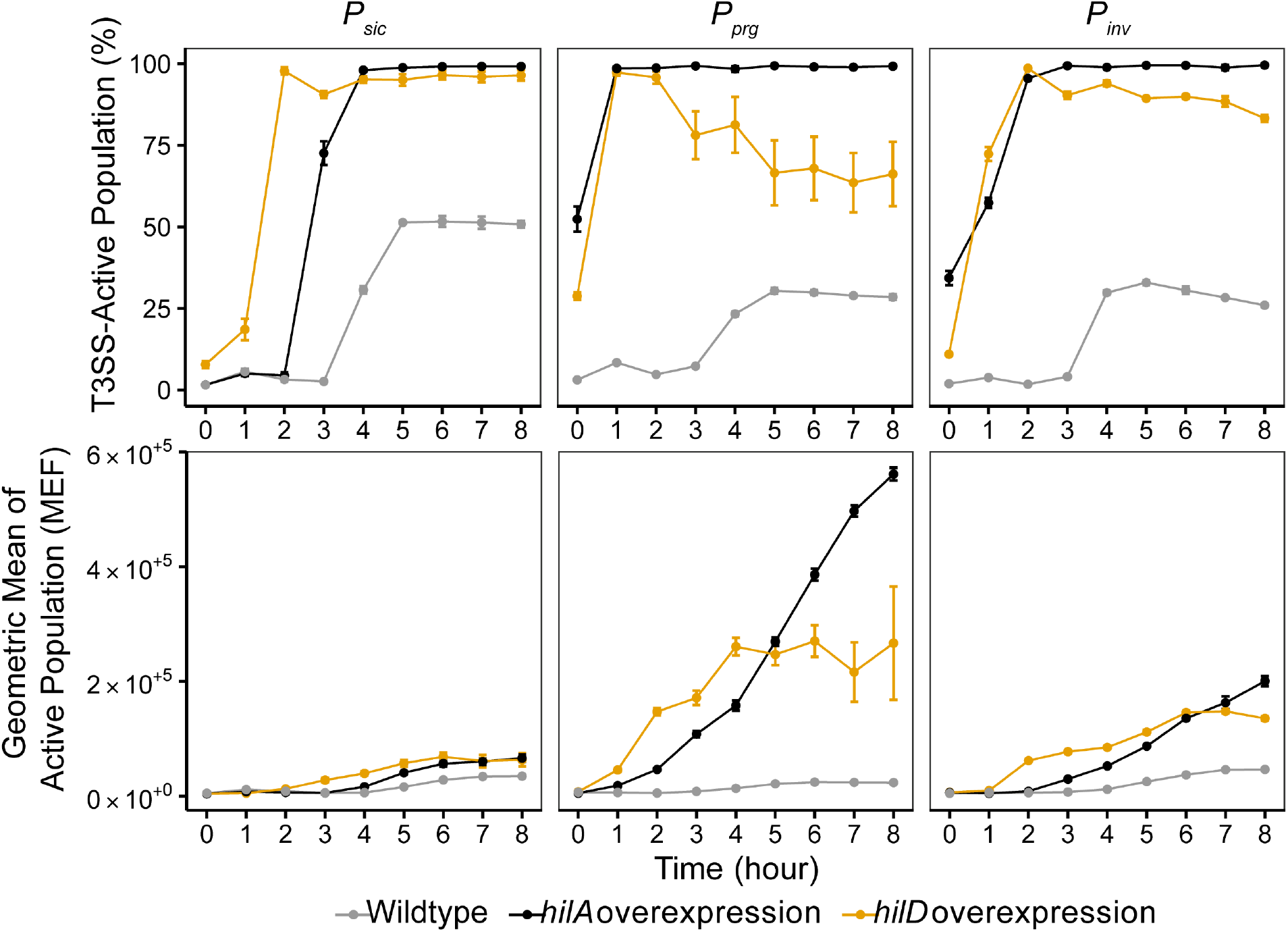
Effect of *hilD* overexpression on transcriptional activity from SPI-1 T3SS operon promoters. Strains contain a plasmid encoding GFP under the control of *P_sic_*, *P_prg_*, or *P_inv_* and either no activation plasmid (wildtype) or an activation plasmid containing either *P_lac_ hilA* or *P_lac_ hilD*. Top: Graph of percentage of population expressing GFP versus time (hours). Bottom: Graph of GFP geometric mean fluorescence intensity (MFI) of the T3SS-active population versus time (hours). GFP fluorescence is representative of T3SS pathway activity from the corresponding promoter. The MFI is representative of the level of expression of the T3SS pathway. Overexpression of *hilA* and *hilD* were induced at 100 μM IPTG. Data represents three replicates and error bars represent standard error.

The *prg* and *inv* operons showed different dynamics than the *sic* operon. Strains carrying either a *hilA* or *hilD* overexpression plasmid showed activity immediately after subculture, while increased activity in the WT strains was delayed until between 3 and 4 hours, similar to the *sic* operon (Figure 3). Surprisingly, both *hilA* and *hilD* overexpression resulted in similar activation dynamics for both *prg* and *inv*. Although both HilA and HilD are known to directly interact with the *inv* promoter, HilD is not known to directly activate *P_prg_* as HilA does.^12, 36–39^ Interestingly, synthetic overexpression of *hilD* caused an early arrest of T3SS activity at both the *prg* and *inv* loci between 2 and 3 hours, as indicated by the drop in the percentage T3SS-active population at these time points. This is different from the activation profile that results from *hilA* overexpression, which shows stable activation of >95% of the population at all time points after 2 hours. Despite the similarity in early activation dynamics between *P_inv_* and *P_prg_*, *hilD* overexpression resulted in early increase and then eventual decrease in GFP levels across all three promoters.

### Genomic modifications that modulate HilD cause constitutive T3SS expression and increase secretion titer

Having established that *hilD* is a promising activator of the T3SS, we next sought to explore non-plasmid-based methods for manipulating *hilD* expression and activation of the T3SS. In doing so, we hoped to improve *hilD*-based activation of the T3SS in two ways. First, we hypothesized that genome-level modifications to *hilD* regulation might eliminate the early deactivation of the T3SS observed in our transcriptional activity data. Second, we wished to eliminate the need for a *hilD* overexpression plasmid to reduce the cost of operating our T3SS secretion system while simultaneously relieving the metabolic burden of sustaining a *hilD* overexpression plasmid. To test non-plasmid-based methods of manipulating *hilD*, we first deleted a gene known to modulate HilD activity as a transcription factor. The SPI-1 T3SS regulator HilE negatively regulates HilD by binding to and inhibiting HilD protein from activating T3SS promoters.^24^ Thus, knocking out *hilE* should have the effect of increasing the amount of available HilD protein, thereby increasing T3SS activation and heterologous protein secretion. Indeed, we found that a strain carrying a deletion of *hilE* (Δ*hilE*) and the export plasmid, *P_sic_* SptP-DH, resulted in secretion titers 2.5-fold higher than a strain without synthetic activation (WT). The Δ*hilE* strain also achieved secretion titers, without synthetic induction, that were 85% of those observed using *hilA* overexpression (Figure 4A).

**Figure 4:**
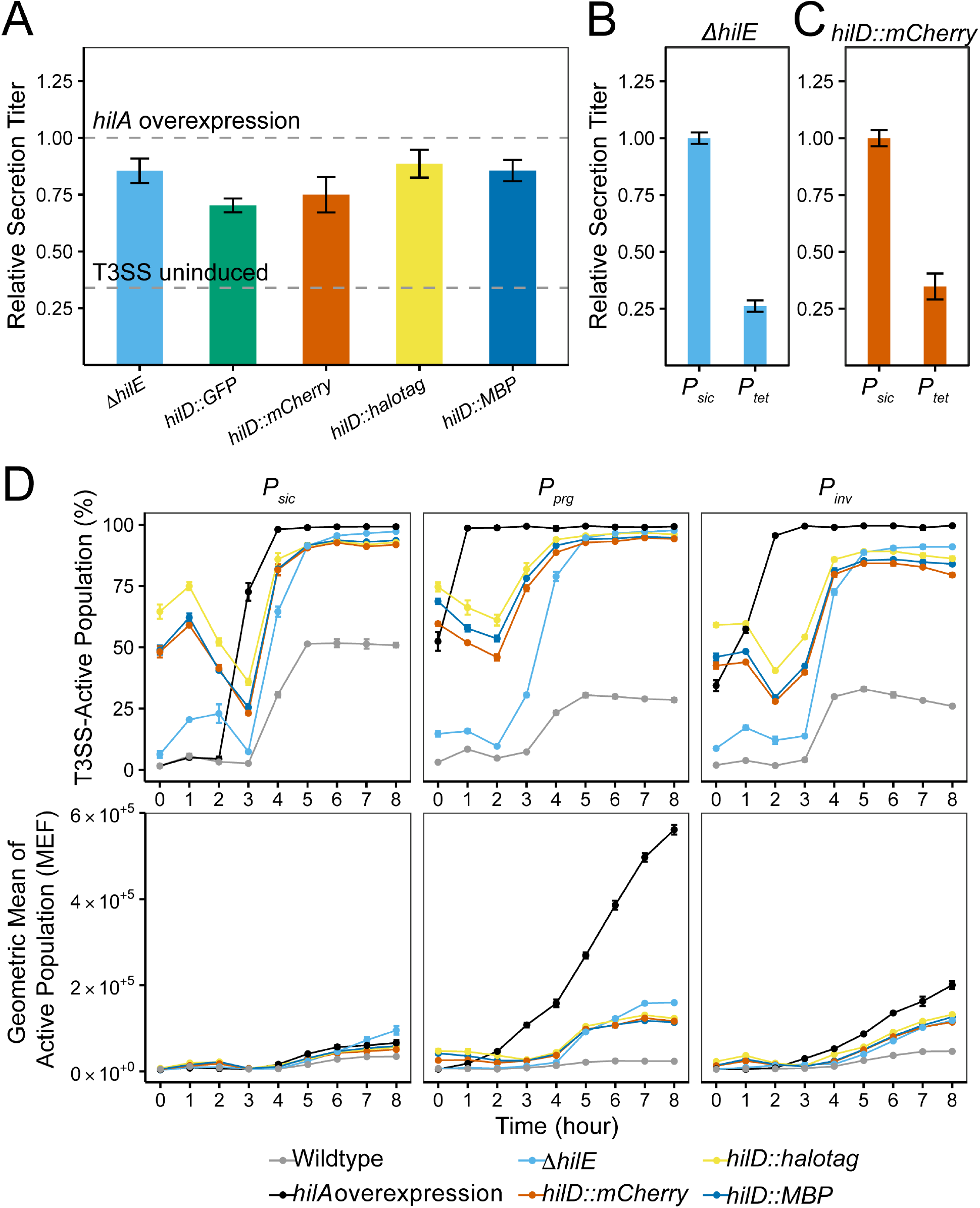
Effect of genomic modifications on protein secretion titer and transcriptional activity from SPI-1 T3SS promoters. **(A)** Relative secretion titer of SptP-DH from genomically modified strains (Δ*hilE*, *hilD::GFP*, *hilD::mCherry*, *hilD::halotag* and *hilD::MBP*) containing the export plasmid encoding *P_sic_* SptP-DH. Secretion titer was measured by densitometry on western blots (probed against an included FLAG tag on the SptP-DH construct) and normalized by OD_600_ and *hilA* overexpression at 100 μM IPTG per replicate (top dashed reference line). The bottom dashed reference line indicates relative SptP-DH secretion titer in a strain without any synthetic induction plasmid. **(B & C)** Comparison of secretion titers when SptP-DH is expressed from the native *sic* promoter versus the *tet* promoter in the **(B)** Δ*hilE* strain and the **(C)** *hilD::mCherry* strain. *P_tet_* SptP-DH is induced with aTc (100 ng/mL) at the optimal induction time for protein secretion (at 4 hours for Δ*hilE* and 3 hours for *hilD::mCherry*). Secretion titer was measured by densitometry on western blots. The titer was normalized by OD_600_ and the average densitometry of *P_sic_* SptP-DH for each activation method. **(D)** Transcriptional activity from strains containing either *P_sic_* GFP, *P_prg_* GFP, or *P_inv_* GFP in wildtype (no synthetic induction), *hilA* overexpression, Δ*hilE*, and the *hilD* transcriptional fusions. Top: Graph of percentage of population expressing GFP versus time (hours). Bottom: Graph of GFP geometric mean fluorescence intensity (MFI) of the T3SS-active population versus time (hours). GFP fluorescence is representative of T3SS pathway activity from the corresponding promoter. The MFI is representative of the level of expression of the T3SS pathway. Overexpression of *hilA* was induced at 100 μM IPTG. In each subfigure, data represents three replicates and error bars represent standard error.

While investigating the effects of plasmid-based *hilD* overexpression and *hilE* knockout on T3SS activation, we constructed fluorescent genomic reporters to monitor *hilD* transcript levels. One iteration of those constructs consisted of GFP or mCherry with its own ribosome binding site inserted immediately downstream of the stop codon of the full-length *hilD* gene. This insertion appeared to confer an increase in SptP-DH titer compared to a WT strain without synthetic *hilA* overexpression. Recognizing that the *hilD* 3’ untranslated region (UTR) functions to regulate *hilD* expression^23^ and insertion of the genes encoding fluorescent reporters shifted the *hilD* 3’UTR further downstream and away from *hilD* (**Supplemental Figure S8**), we created two additional strains with the common reporter proteins HaloTag and maltose-binding protein (MBP) inserted similarly to determine if the effect was general. All *hilD* transcriptional fusions produced secretion titers above the uninduced WT and between 70-89% of that produced by *hilA* overexpression (Figure 4C). Expression levels of SptP-DH for all strains except *hilD::GFP* were 1.2-1.5-fold higher than that produced by *hilA* overexpression (**Supplemental Figure S9**). This is similar to the increase in protein expression seen when the T3SS is activated by *hilD* overexpression (**Supplemental Figure S7**). This suggests that the genomic modifications explored herein may be activating the T3SS through a similar route as *hilD* overexpression.

We next investigated the impact of Δ*hilE* and the *hilD* transcriptional fusions on T3SS activation. We measured T3SS activation using the same flow cytometry assay described earlier, in which the readout is GFP expression under the control of T3SS promoters *sic*, *prg*, and *inv*. We did not include *hilD::GFP* because it would interfere with the plasmid-based GFP reporter on T3SS promoter activity. The Δ*hilE* and the *hilD* transcriptional fusions show different T3SS transcriptional dynamics compared to WT without induction and the synthetically activated strains (Figure 4C). The strains containing the *hilD* transcriptional fusions (*hilD::mCherry*, *hilD::halotag*, and *hilD::MBP*) show constitutive activation from the overnight culture for all three promoters, indicated by the 42-75% active population observed for these strains immediately at subculture (0 hours on the plot). These same strains then undergo subsequent relaxation, indicated by a drop in the active population at 2 hours for *prg* and *inv* and 3 hours for *sic*. Finally, the increase in percent active population shows that these strains eventually reactivate after this relaxation.

The Δ*hilE* strain, however, had only a small active population from the overnight culture (6-15% active population across promoters). The Δ*hilE* strain lacked a relaxation and showed an activation at 2 hours for *prg* and 3 hours for *inv* and *sic*. Despite the different activation dynamics, all four strains reach similar amounts of active populations (>92%, >94%, >79% for *sic*, *prg*, and *inv* respectively) and reach similar levels of mean fluorescence in the active population at the *inv* and *prg* loci (Figure 4C). The Δ*hilE* strain activated a higher percentage of the population for all loci (97%, 98%, 91% for *sic*, *prg*, and *inv* respectively) and showed a 1.6-fold higher maximum MFI at the *sic* operon than the highest *hilD* transcriptional fusion, *hilD::MBP*. These data show that there are differences in transcriptional activity between the Δ*hilE* and the *hilD* transcriptional fusion strains even though both strain sets exhibit similar secretion titers and elevated POI expression. This suggests that Δ*hilE* and the *hilD* transcriptional fusions likely regulate *hilD* in different ways. The *hilE* knockout modulates at the translational level^24^, whereas we hypothesize that the *hilD* transcriptional fusions modulate *hilD* at the post-translational levels. Importantly, all the genomic modifications explored avoid the early arrest in T3SS activation that occurs when *hilD* is overexpressed from a plasmid.

Next, we wanted to confirm that the *sic* promoter retained its ability to adjust for optimal POI expression with changing T3SS induction strategies. We determined the optimal induction times to induce *P_tet_* DH in the Δ*hilE* and *hilD::mCherry* strains (**Supplemental Figure S10**) and compared secretion titers between *P_sic_*- and *P_tet_*-driven expression of the POI. The *P_sic_*-driven expression resulted in a statistically significant (p-value < 0.05) 3.8- and 2.9-fold higher SptP-DH secretion titers for Δ*hilE* and *hilD::mCherry* compared to *P_tet_* (Figure 4B).

### Genomic modifications combine to make constitutive SPI-1 T3SS strain with comparable secretion titers to plasmid-based activation strategies

The Δ*hilE* strain and the *hilD* transcriptional fusions resulted in constitutive expression of the T3SS but have different theoretical mechanisms for manipulating HilD and different transcriptional dynamics. Therefore, we explored whether combining the different mutations might increase secretion titers further. We constructed a double mutant strain with Δ*hilE* and one of the *hilD* transcriptional fusions. Noting that all the transcriptional fusions behaved similarly with respect to secretion titer and transcriptional activity, we moved forward with just one of the fusions, mCherry, in the double mutant strain. We chose mCherry because it could serve as a useful cellular marker as a fluorescent protein in future studies. We measured SptP-DH secretion titer in the Δ*hilE hilD::mCherry* double mutant strain and compared it to the single mutants and to *hilA* overexpression. The Δ*hilE hilD::mCherry* strain secreted SptP-DH at levels equivalent to *hilA* overexpression, exceeding the secretion titers achieved with each individual genomic modification (Figure 5). The double mutant had a higher constitutive activation in the overnight culture than the individual *hilD* transcriptional fusions, ranging from 92%-96% active across loci (**Supplemental Figure S14**). It also showed the same relaxation and then reactivation dynamics as observed for the individual *hilD* transcriptional fusions at each locus. After 8 hours, almost 100% of the population was T3SS-active. This suggests that the increased secretion titer in the Δ*hilE hilD::mCherry* strain may be due to the increased T3SS-activated population. Most importantly, the double mutant strain successfully bypasses the early deactivation of the T3SS that we see when *hilD* is synthetically overexpressed. When we compared *P_sic_*- and *P_tet_*-driven secretion titers, the *sic* promoter had a statistically significant 1.6-fold higher secretion titer compared to *P_tet_* (p-value < 0.05, **Supplemental Figure S16**). This confirms that *P_sic_* is still appropriate to use to drive expression of the POI.

**Figure 5:**
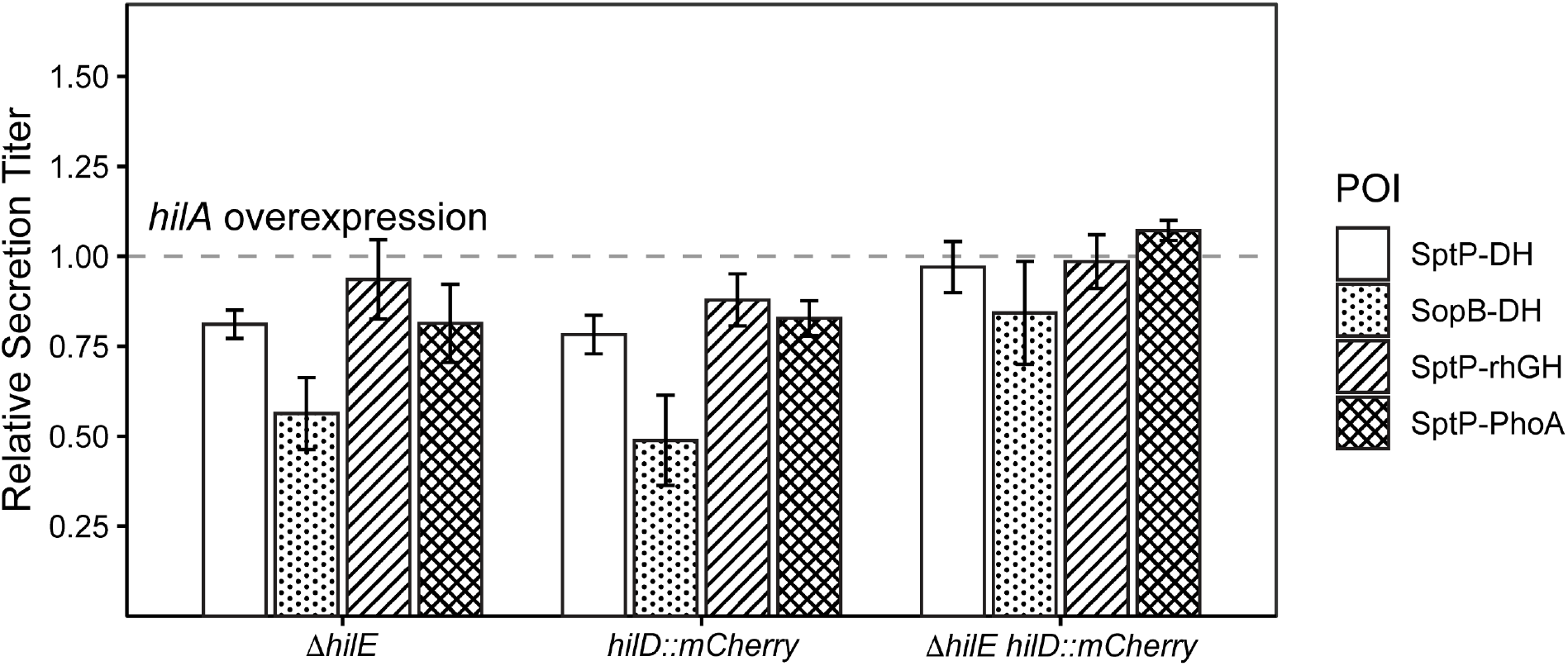
The genomic modifications combine to make a constitutively active T3SS that secretes a variety of proteins. Relative secretion titer of a range of proteins (SptP-DH, SopB-DH, SptP-rhGH, SptP-PhoA) under the control of *P_sic_* and secreted by the Δ*hilE*, *hilD::mCherry*, and the Δ*hilE hilD::mCherry* double mutant strains. Protein expression was measured by densitometry on western blots (probed against a FLAG tag included in each construct) and normalized by OD_600_ and *hilA* overexpression at 100 μM IPTG per replicate (dashed reference line). Data represents three replicates and error bars represent standard error.

Finally, we evaluated whether the effect of the constitutive Δ*hilE hilD::mCherry* strain was universal in secreting other POIs or when using a different secretion tag. First, we chose SopB^9^ as an alternative secretion tag, replacing the SptP tag in the original export fusion. We also selected human growth hormone (mature somatropin, rhGH) and alkaline phosphatase (PhoA) as alternative secreted proteins, replacing the DH protein in the original fusion.^6, 20^ We measured the expression and secretion titer of SopB-DH, SptP-rhGH, and SptP-PhoA in the Δ*hilE, hilD::mCherry*, and Δ*hilE hilD::mCherry* strains. Results were normalized to the secretion titer using *hilA* overexpression. As a general trend, the Δ*hilE hilD::mCherry* strain secretes at levels close to or higher than strains that activate the T3SS by overexpressing *hilA*. For example, when expressing SopB-DH, the Δ*hilE* strain and *hilD::mCherry* strain secreted at levels 57% and 50% of levels observed upon *hilA* overexpression, respectively (Figure 5). In contrast, the secretion level of SopB-DH in the double mutant strain, Δ*hilE hilD::mCherry*, was not statistically different from secretion levels in a strain that activated the T3SS via *hilA* overexpression. Likewise, the double mutant strain was capable of secreting SptP-rhGH and SptP-PhoA at titers equivalent to those produced by a strain that activates the T3SS via *hilA* overexpression. These data show that the two genomic modifications (Δ*hilE* strain and *hilD::mCherry*) combine to make a constitutively active T3SS capable of secreting a range of POIs tagged with different signal tags while avoiding the early deactivation that comes with *hilD* overexpression.

## Discussion

We set out to identify new ways to activate the *S*. *enterica* SPI-1 T3SS to achieve high secretion titers for heterologous protein production. In the process, we made several unexpected but important observations related to the regulation of the system. These observations include differences between regulators *hilA* and *hilD*, the continued utility of the *sic* promoter in driving POI expression, and a role for the *hilD* 3’UTR in T3SS heterologous protein secretion.

We noticed differences in transcriptional activation profiles between *hilA* and *hilD* overexpression as T3SS activation methods. For example, when we activated the T3SS using *hilD* overexpression, we were surprised to see the early activation of the *sic* operon because *hilD* is upstream of *hilA* in the regulatory cascade and we thus expected that activation by *hilD* would result in later activation. HilD is not known to activate the *sic* promoter directly, but instead indirectly through the *inv* operon.^40^ However, both *hilA* and *hilD* overexpression activated the *inv* operon at the same time and we do not see the earlier *sic* operon activation with *hilA* overexpression. One possible explanation is that overexpression of *hilD* overcomes any repressing mechanisms on the SPI-1 T3SS promoters, resulting in earlier activation of the *sic* operon. HilD activates *hilA*,^15^ its own promoter, and the promoters of other T3SS transcriptional regulators.^41^ This results in a feed-forward loop that accelerates SPI-1 T3SS expression once a threshold HilD level is exceeded. Interestingly, earlier activation of the T3SS did not result in statistically significantly higher secretion titers per cell. This may be because *hilD* overexpression led to lower cell densities and an early arrest in *sic*, *prg*, and *inv* expression, suggesting that too much HilD hinders other factors that impact secretion titers.

The *sic* promoter consistently works well for driving expression of our POI, regardless of the T3SS activation method. Use of the *tet* promoter to drive POI expression requires re-optimization of the induction timing and level with each new activation method. Although the *tet* promoter resulted in similar secretion titers to the *sic* promoter in one case (Figure 2B), it resulted in lower secretion titers in other cases (Figure 2A, Figure 4BC, **Supplemental Figure S16A**). This is in agreement with the observation from the T3SS transcriptional data that T3SS regulation is controlled differently between activation methods. In contrast, the *sic* promoter consistently resulted in similar or higher secretion titers than the *tet* promoter. We believe the use of the *sic* promoter has some additional effect that enables a balance between expression and secretion. For example, the Δ*hilE* strain resulted in significantly higher secretion titers with the *sic* promoter than the *tet* promoter, which is partially explained by the differences in expression level (Figure 4B, **Supplemental Figure S12A**). However, the *hilD::mCherry* strain gave rise to significantly higher secretion titers with the *sic* promoter than the *tet* promoter while conferring similar expression levels of the POI between the two promoters (Figure 4C, **Supplemental Figure S12B**). This resulting difference could be because we only optimized the *tet* promoter by hour intervals, and to match the *sic* promoter in secretion titers requires more fine-tuned optimization using shorter time intervals. Alternatively, the *sic* promoter may confer higher secretion titers because it enables dynamic expression of the T3SS pathway over time, whereas the *tet* promoter provides uniform gene expression over time once induced. Regardless of the method of T3SS activation, the *sic* promoter confers optimal expression of the POI for secretion. This is advantageous because it successfully bypasses the optimization steps that an inducible promoter requires. It also means that we do not have to induce the POI separately from activation of the T3SS, reducing the complexity of our protein secretion system. Altogether, these data support the continued use of the native *sic* promoter to express the POI when using the T3SS for protein secretion.

While exploring methods to modulate *hilD* activation of the T3SS, we found that the *hilD* 3’UTR presented an orthogonal handle to activate the system and opened the door to a new engineering route. Both the deletion of *hilE* and the *hilD* transcriptional fusions resulted in constitutive T3SS activity and increased protein secretion titer. While this result was expected for Δ*hilE*, we did not anticipate that *hilD* transcriptional fusions would result in such an increase. We propose that the added DNA downstream of the *hilD* coding sequence contributes to a stabilizing effect of the mRNA encoding for *hilD*. The *hilD* 3’UTR negatively regulates *hilD* expression,^23^ so the added DNA may increase HilD protein levels by stabilizing the mRNA and contributing to ribosome shielding of the *hilD* transcript.^42^ More stable *hilD* mRNA may also have other regulatory impacts, because the *hilD* 5’UTR contains stem loops that modulate invasion gene expression.^43^ Although we did not intentionally engineer the 3’UTR to control *hilD* expression, modifying the 3’UTR could be a complementary genome engineering approach to changing the promoter to control gene expression, especially for bacterial hosts. Fine-tuning expression of target genes by engineering the 3’UTR can be particularly beneficial in situations like those described here, in which the plasmid overexpression firehose approach is not ideal.

Finally, we constructed a constitutively active T3SS strain, Δ*hilE hilD::mCherry*, that successfully secreted a variety of different POIs with different signal tags. Since IPTG is an expensive inducer to use (21% - 28% of the total media cost, for industrial and academic uses, respectively, Table 1), this new strain secretes at levels comparable to *hilA* overexpression without the need for an added inducer. The constitutively active strain also has one less plasmid which reduces the complexity and chance for plasmid instability, making it more suitable for industrial fermentations. This lowers the metabolic load imposed by plasmid maintenance, increases stability of the system over time, and decreases the overall cost of protein production. We expect that exploring analogous regulatory strategies in other contexts may prove useful not only in controlling a variety of other technologically relevant biological systems, but also in gaining more insight into underlying regulatory network behavior, as we observed here.

## Acknowledgements

We want to acknowledge and thank everyone in the Tullman-Ercek lab who supported us and made publication of this paper possible. Special thanks to Dr. Carolyn Mills, Samuel Leach, and Nolan Kennedy for their invaluable feedback throughout the writing of this manuscript. We would also like to thank Dr. Kevin J. Metcalf for the original construction of *P_prg_* GFPmut2.

JML was supported by the National Institutes of Health Training Grant (T32 GM008382) through Northwestern University’s Molecular Biophysics Training Program. HTW was supported by the National Science Scholarship from the Agency of Science, Technology and Research, Singapore. LAB was supported by a National Science Foundation Graduate Research Fellowship. DTE was supported by the National Science Foundation (award number BBE-1706125 to DTE).

